# resolveS: rapid inference of RNA-seq library strandedness using universal rRNA alignments

**DOI:** 10.64898/2026.01.08.698333

**Authors:** Xi Long, Ting Zhao, Dalang Yu

## Abstract

Accurate specification of RNA-seq library strandedness is essential for read counting, transcript assembly and antisense transcription analysis, yet this information is frequently missing from public metadata. We present resolveS, a fast and lightweight method that infers strandedness from alignments to a compact universal rRNA database, avoiding the need for organism-specific reference genomes and annotations. Across benchmark datasets, resolveS provides robust strandedness calls while maintaining very low runtime and memory usage, making it suitable for both routine quality control and large-scale reanalysis of public RNA-seq data.

## Introduction

High-throughput RNA sequencing (RNA-seq) has revolutionized our understanding of transcriptomes, enabling precise quantification of gene expression, alternative splicing, and novel transcript discovery (Conesa, et al., 2016; Gondane and Itkonen, 2023; Hong, et al., 2020; Sahraeian, et al., 2017; Stark, et al., 2019). Modern RNA-seq protocols frequently employ strand-specific library preparation (Figure 1A), a capability essential for distinguishing transcripts originating from overlapping genes encoded on opposite strands. The overlapping genes compose approximately a quarter of human protein-coding genes and are widespread in compact genomes (Chen, et al., 2019). Incorrect specification of the strandedness parameter in downstream analysis tools (Figure 1B), such as featureCounts (Liao, et al., 2014) or Trinity (Haas, et al., 2013), can severely bias analytical results. Despite the importance of this this parameter, it is frequently omitted from the metadata in public repositories, such as the European Nucleotide Archive and Sequence Read Archive, and is often absent from corresponding publications. Notably, a systematic investigation found that approximately 44% of associated studies failed to explicitly report the library preparation method(Signal and Kahlke, 2022).

**Figure 1.**
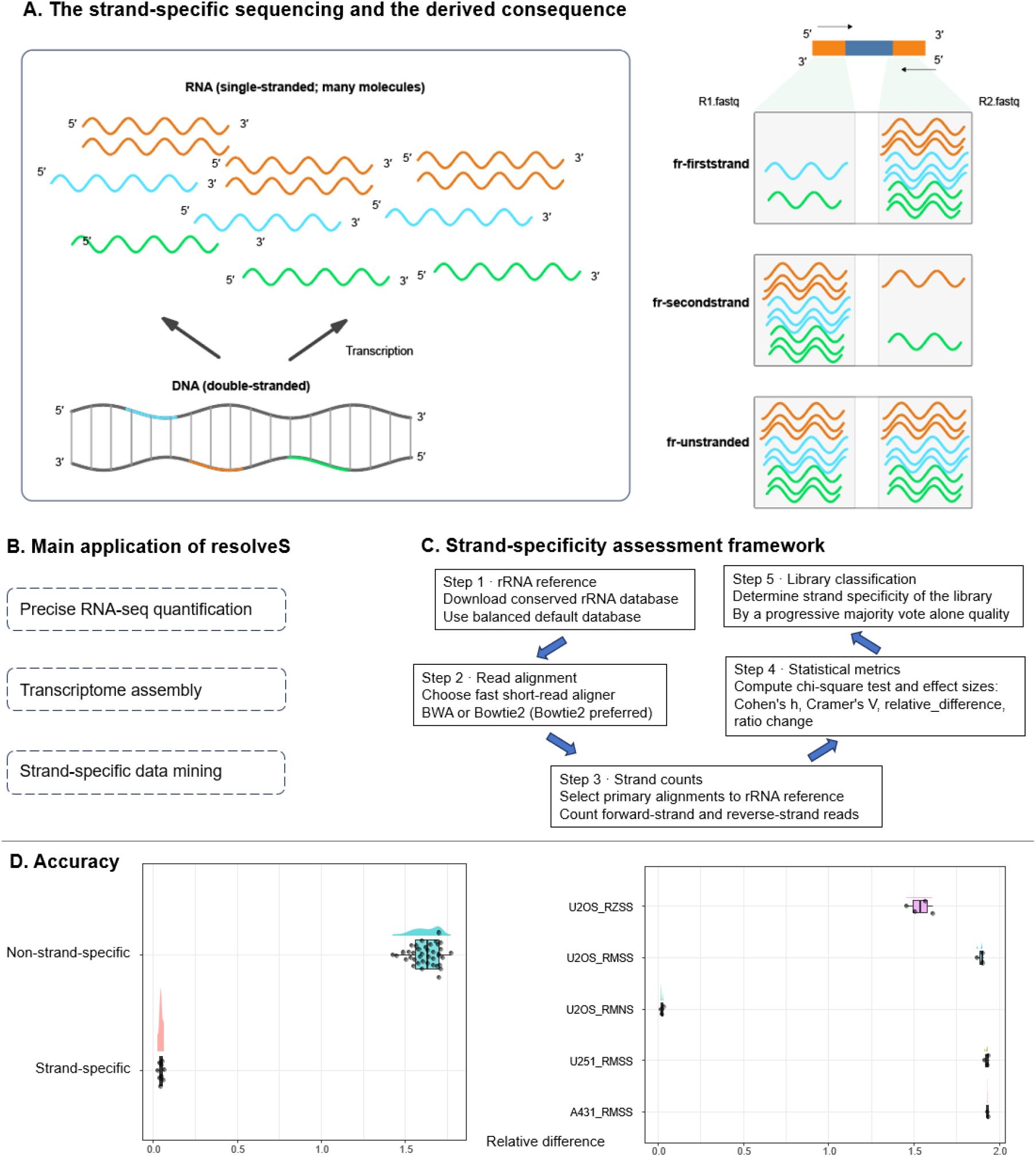
Accuracy, robust framework and application of resolveS. A. The principle of the resolveS. The strandness of the RNA is defined by the RNA molecule and the derived consequences is intuitional for the “fr-firststrand” type is R2 sequence same as the original RNA and the “fr-secondstrand” is vice versa. For the authenticity, a small fraction of the RNA may contain the Nature Antisense transcripts. B. Overview of the main application areas of resloveS. C. The framework summary: First, A conservative, universal rRNA database with default version (https://github.com/sortmerna/sortmerna/releases) is downloaded and indexed. Next, reads alignment with the curated maximum size sampling using Bowtie2. Third, only primary alignments are retained, and counts for forward and reverse strands are recorded. Then, Strandedness Inference: Several statistics are computed. We put forward several non-sample size aware statistics as indices to estimate the strandedness. Finally, Library classification by the statistics. D. Accuracy and robustness of the resolveS. The representative statistics (Relative difference) have the maximum range to distinguish the strandedness. [∗RM = enriched with RiboMinus, RZ = enriched with RiboZero, SS = strand specific, NS = non-stranded. A431, U251 and U2OS denote different cell lines] (Sigurgeirsson, et al., 2014). Additional statistical distributions and full details are provided in the supplements.

Existing tools attempt to address this demand but face significant limitations. For instance. RSeQC’s infer_experiment.py requires a genome-aligned BAM file and genome annotation file recodes the gene information with bed format (Wang, et al., 2012). Similarly, how_are_we_stranded_here (Signal and Kahlke, 2022) improves speed by using pseudoalignment on a subset of reads to generate a genome-sorted BAM file before running infer_experiment.py. However, a fundamental limitation of these methods is their reliance on species-specific reference genomes and GTF annotation files. This dependence poses a significant challenge for studies involving newly sequenced or non-model organisms where high-quality genomic resources are unavailable. It is worthy note that there is complex pipeline to address this problem, one can use the Trinity (Haas, et al., 2013) to assemble the transcriptome data first (with guess of 3 types of the strandedness Figure 1A) and then estimate the annotation file via the TransDecoder (https://github.com/TransDecoder/TransDecoder) and convert the gff3 to bed file by BEDOPS (Neph, et al., 2012), next align the fastq to the assembly to get the bam file, finally employ the RSeQC to infer the strandedness by input the gff3 and bed. The solution takes user to install plenty of software and operates several procedures which hinds the large-scale application. Furthermore, the accuracy and validity of this ad hoc strategy have not been rigorously established.

In this work, we developed resolveS, an ultra-fast, reference-agnostic solution. The principal novelty of resolveS lies in its independence of species-specific genomics annotations, instead, it utilizes a compact, curated database of universal rRNA sequences as a mapping target. This design leverages the evolutionary conservation of ribosomal RNA (rRNA) across the tree of life and the abundant known knowledge making them an ideal universal proxy. Furthermore, we address a common statistical pitfall in NGS data analysis—the sensitivity of P-values to large sample sizes—by implementing a progressive detection strategy that evaluates rRNA targets based on mapping quality. In addition to reporting the strandedness type, resolveS also provides confidence estimates under various complex scenarios. By combination this universal rRNA database with the optimized, stop-on-max-reads functionality of Bowtie2<Langmead, 2012 #16}, resolveS achieve remarkable speed and resource efficiency, establishing it as a suitable tool for universal strandedness-detection.

## Materials and methods

### The test datasets

Data sets with ground truth from the animal and plants(Shi, et al., 2025) are tested. The animal cell line data is from our own unpublished and others from Naphade (Naphade, et al., 2022), Sigurgeirsson (Sigurgeirsson, et al., 2014). Large scale benchmarking data from Signal(Signal and Kahlke, 2022) supplement collection. Five samples from the Signal et al. are removed, according to careful evaluation.

### The universal rRNA reference strategy

We employed a lightweight, universal reference database that derived from the SortMeRNA database (Kopylova, et al., 2012). which includes representative rRNA sequences. There are three versions of the rRNA files, we choose the default version as default reference based on our empirical experiencing and practical comparisons, while a high-sensitivity version is provided as an alternative.

Details indexing instructions are available at https://github.com/yudalang3/resolveS/tree/main/docs.

### Aligner choice and subsampling optimization strategy

Speed and memory efficiency were primary design goals. We compared established DNA aligners, form matured BWA(0.7.19-r1273) (Li and Durbin, 2009), Bowtie2 (2.5.4) (Langmead and Salzberg, 2012) to the fresh LexiMap(v0.8.0) (Shen, et al., 2025) and found Bowtie2 is best for our assignment because it significantly outperforms BWA and LexiMap for this specific task:

Speed Optimization: Bowtie2 supports the -u parameter to stop alignment after a specified maximum number of reads, dramatically accelerating the inference process. BWA lacks this feature. The LexiMap is not pretty well for mapping the fastq millons numbers of reads.

Resource Efficiency: Bowtie2’s index files are smaller (214 MB vs. 521 MB for BWA), and the original FASTA library file is not required after indexing, reducing storage overhead. Furthermore, testing showed Bowtie2 requires half the memory and runs 2-3 times faster than BWA for alignment.

After evaluated such mainstream aligners, we selected Bowtie2 for its outstanding performance of speed and low memory footprint.

Unlike tools that map the entire FASTQ file, resolveS uses an efficient early-stopping subsampling strategy that limits alignment to the first batch of reads. We utilize the -u (stop after N reads) capability of Bowtie2 to map only the first sufficient batch of reads (default: 1,000,000 reads) to the rRNA index. This reduces the runtime complexity from O(N) (where N is total library size) to O(1) (constant time), allowing the tool to return results in seconds regardless of whether the input file is 100 MB or 100 GB.

### Statistical Framework for Strandedness Inference

Besides the commonly used Chi-square test. Several simple but powerful effect size metrics (“Fwd_Ratio”, “F2R_Ratio”, “Rel_Diff”, “Cohens_h”, “Cramers_V”, “Bayes_Factor”, “Epsilon”, “Hellinger”, “Entropy”) were implemented to provide robust inference of the strandedness, see https://github.com/yudalang3/resolveS/blob/main/check_strand.py for details.

We selected the signed symmetric relative difference (Rel_Diff) as the primary decision statistic because it provided the clearest separation between stranded and unstranded libraries in benchmark data. We choose the empirical value abs (Ref_Diff) = 0.6 as the cutoff for the stranded-specificity and the number of samples for the “insufficient-data” to be less than 18. The rationality is validated by the test datasets.

### The strandedeness detection level

Earlier versions inferred strandness from global fragment counts, whereas the current implementation determines strandness via per□chromosome voting among the most covered chromosomes (with adaptive MAPQ filtering). The “resolveS_fast” version still employs this approach.

Strandedness detection uses a progressive majority vote across the top chromosomes ranked by valid read counts, but it is run under an adaptive MAPQ filter. The algorithm first evaluates reads with MAPQ ≥ 20; if the detection level is all-insufficient-fallback, it retries with lower thresholds (MAPQ 10, then 3, then 0) until sufficient evidence is found or all levels fail. For each run, chromosomes with too few reads are labeled “insufficient-data” and excluded from voting. The majority check proceeds 3/3 → 4/5 → 6/7 → 7/8, and the first passing level is reported (e.g., 3of3, 4of5, 6of7, 7of8). If no majority is reached, a fallback level is recorded and a global forward/reverse ratio is calculated to determine the final inference.

## Results

### An accuracy- and robustness-oriented framework for strandedness detection

The resolveS workflow is streamlined as a dedicated pipeline for maximum efficiency (Figure 1C, Table 1). After tests several aligners, we found the Bowtie2 is best suit for our purpose (Method section2). We tested the rRNA database version provided by the SortMeRNA, the default version is almost no difference with the sensitive version. The input RNA volume and RNA enrichment experiment methods even the RiboZero protocol is not confounding the accuracy (Figure 1D). Although RNA-seq library preparation protocols, such as poly(A) enrichment or Ribo-Zero depletion, aim to remove rRNA, these methods are rarely 100% efficient. resolveS exploits residual rRNA reads as an informative signal for strandedness inference.

**Table 1.** The summary of the running speed and memory usage of the resolveS_fast.

| Input type | Approach | Threads number | Maximum align reads | Time cost (s) | Memory cost (GiB) |
| --- | --- | --- | --- | --- | --- |
| Batch_gzip_fastq_5 | Apptainer_image | 15 | 1000000 | 18.7733(0.3099) | 0.3240(0.0017) |
| Batch_gzip_fastq_5 | Apptainer_image | 15 | 4000000 | 68.0333(4.3625) | 0.3277(0.0025) |
| Batch_gzip_fastq_5 | Apptainer_image | 8 | 1000000 | 21.1567(0.4267) | 0.2813(0.0021) |
| Batch_gzip_fastq_5 | Apptainer_image | 8 | 4000000 | 75.1567(2.7356) | 0.2833(0.0006) |
| Batch_gzip_fastq_5 | Bash_script | 15 | 1000000 | 21.9400(0.5011) | 0.3280(0.0017) |
| Batch_gzip_fastq_5 | Bash_script | 15 | 4000000 | 79.6300(2.5363) | 0.3310(0.0000) |
| Batch_gzip_fastq_5 | Bash_script | 8 | 1000000 | 19.8900(0.5862) | 0.2833(0.0021) |
| Batch_gzip_fastq_5 | Bash_script | 8 | 4000000 | 72.6667(4.3824) | 0.2857(0.0006) |
| Batch_gzip_fastq_5 | Singularity_image | 15 | 1000000 | 19.9367(0.1464) | 0.3267(0.0012) |
| Batch_gzip_fastq_5 | Singularity_image | 15 | 4000000 | 65.6567(0.4693) | 0.3293(0.0006) |
| Batch_gzip_fastq_5 | Singularity_image | 8 | 1000000 | 21.7067(0.5609) | 0.2800(0.0017) |
| Batch_gzip_fastq_5 | Singularity_image | 8 | 4000000 | 70.7333(0.9570) | 0.2857(0.0029) |
| Single_gzip_fastq | Apptainer_image | 15 | 1000000 | 3.6867(0.0252) | 0.3227(0.0006) |
| Single_gzip_fastq | Apptainer_image | 15 | 4000000 | 10.8867(0.2371) | 0.3250(0.0017) |
| Single_gzip_fastq | Apptainer_image | 8 | 1000000 | 4.1033(0.1159) | 0.2783(0.0006) |
| Single_gzip_fastq | Apptainer_image | 8 | 4000000 | 11.7100(0.2706) | 0.2817(0.0023) |
| Single_gzip_fastq | Bash_script | 15 | 1000000 | 2.9900(0.0300) | 0.3243(0.0006) |
| Single_gzip_fastq | Bash_script | 15 | 4000000 | 9.7700(0.6791) | 0.3283(0.0029) |
| Single_gzip_fastq | Bash_script | 8 | 1000000 | 2.9767(0.0231) | 0.2810(0.0000) |
| Single_gzip_fastq | Bash_script | 8 | 4000000 | 10.7567(0.6309) | 0.2850(0.0000) |
| Single_gzip_fastq | Singularity_image | 15 | 1000000 | 3.6133(0.1041) | 0.3240(0.0026) |
| Single_gzip_fastq | Singularity_image | 15 | 4000000 | 11.5267(0.8363) | 0.3283(0.0051) |
| Single_gzip_fastq | Singularity_image | 8 | 1000000 | 3.8433(0.0379) | 0.2807(0.0023) |
| Single_gzip_fastq | Singularity_image | 8 | 4000000 | 11.5700(0.1609) | 0.2790(0.0000) |

The CPU is 13th Gen Intel(R) Core (TM) i7-13700 and storage is WD Green SN350 1TB
The output style is Mean (Standard Deviation)

A common challenge in analyzing NGS count data is the “P-value fallacy”, where trivial deviations become statistically significant simply due to the massive number of reads (counts). A standard Chi-square test might reject the null hypothesis of uniformity even when the deviation is biologically irrelevant (e.g., 50.1% vs 49.9%). To overcome this, we implemented robust, sample-size-independent statistical metrics to ensure reliable inference. All 9 metrices (Methods section 4) are distinguishable to the strand specificity (Supplement Figure1-3).

The relative difference index has the greatest range to distinguish strandedness (Figure 1D). Also, for the simplicity, we reduce reversal metrices to only the relative difference. Besides give final strandedness, we also provide the confident level of the inference. The most reliable level is the “MAPQ-20” and “3of3” means filter the mapping quality by the score 20 and the top 3 rRNA sequence concordantly agree to be one type. While the worst level is the “MAPQ-0” with “multi-of8-fallback”, which means although setting the mapping quality to lowest value, the top 8 rRNA transcripts still give over 2 candidates.

To validate accuracy, we tested resolveS on a ground-truth dataset where library preparation methods were known. Without require any data clean approach, resolveS achieved 98.81% (249/252) concordance with the metadata labels for both unstranded and stranded libraries. And in the failed 3 samples, resolveS correctly gives the clues of the fallback detection level. When compared with existing approaches, resolveS demonstrates superior convenience and efficiency (Table 2).

**Table 2.**
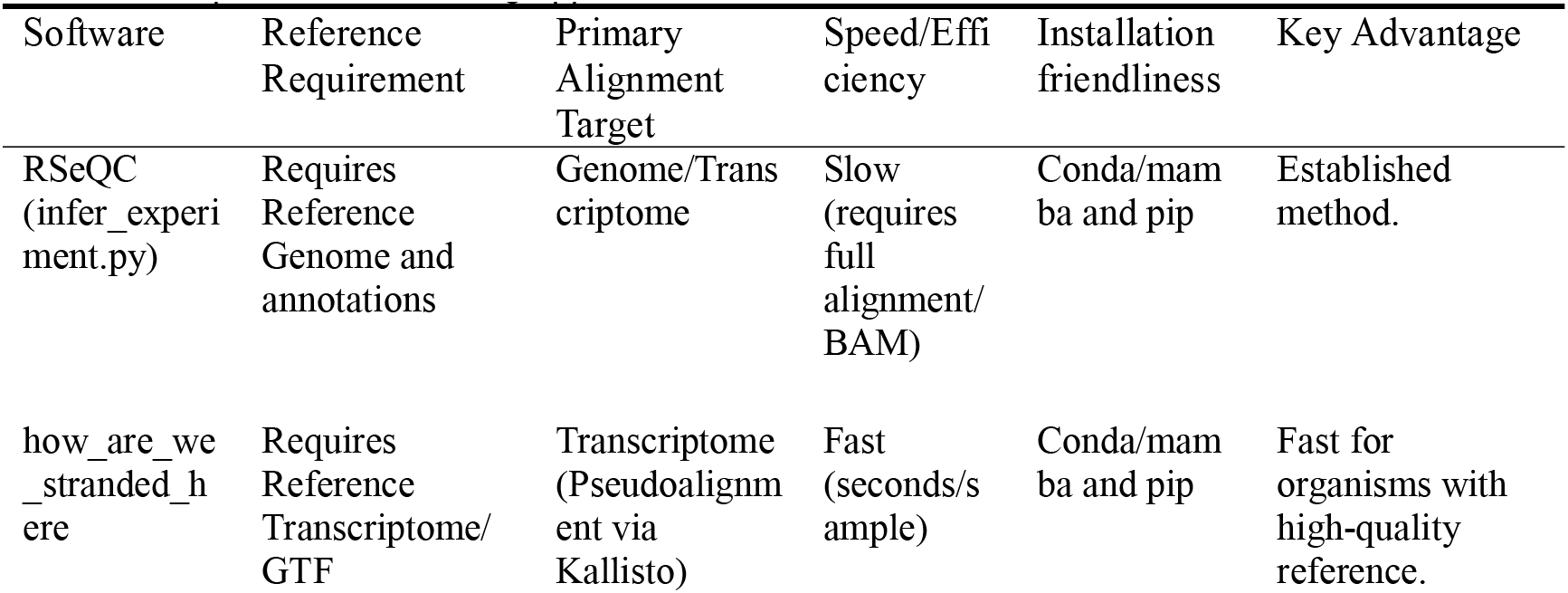
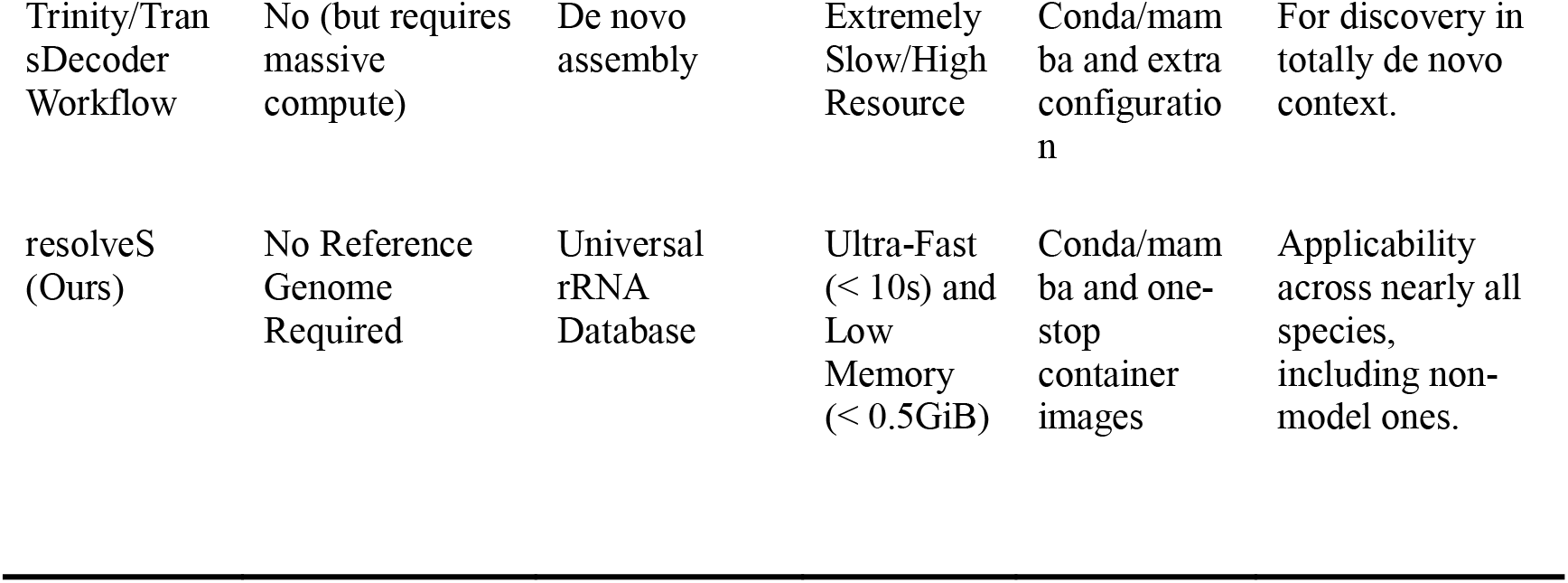
Comparison with Existing approaches to infer strandedness.

### Installation and documentation

All source codes publicly available without restriction at: https://github.com/yudalang3/resolveS. Portable version and container image files can be download directly from the Release page.

### Conclusion and future work

In this work, we present resolveS, an open-source, efficient and user-friendly tool for rapid determining RNA-seq library strandedness and give the detection confident level. FASTQ files from any processing stage (raw or trimmed) are supported, however, we recommend directly using the raw data for simplicity. Its capacity to operate independently of species-specific genomics annotation extends its utility beyond model organisms, ensuring that researchers analyzing non-model or poorly annotated species can still reliably ascertain this critical experimental parameter. We strongly recommend incorporating resolveS into upstream quality control pipelines to ensure the accuracy and reproducibility of downstream quantitative and assembly analyses.

One limitation of current approach is the theoretical dependence on residual rRNA. However, in our extensive testing across samples, we have not encountered datasets with zero rRNA alignments, although ambiguous fallback cases are occasionally observed, most likely due to low-level contamination from other samples. Future work will focus on accelerating the sequencing mapping speed to reduce the total running time.

Specifically, we aim to develop a customized mapping strategy to optimize this task.

## Supporting information

Supplemental Matrial

## Availability and Implementation

resolveS is available under the MIT license at (https://github.com/yudalang3/resolveS). Ready to use singularity and apptainer images as well as portable version available in the release page (https://github.com/yudalang3/resolveS/releases)

## Author contributions

Dalang Yu (Conceptualization [lead], Data curation [lead], Formal analysis [lead], Funding acquisition [lead], Investigation [lead], Methodology [lead], Project administration [lead], Resources [lead], Software [lead], Supervision [lead], Validation [lead], Visualization [lead], Writing—original draft [lead], Writing—review & editing [lead])

Xi Long (Data curation [lead], Formal analysis [lead], Funding acquisition [lead], Investigation [lead], Methodology [lead], Project administration [lead], Resources [lead], Software [lead], Supervision [lead], Validation [lead], Visualization [lead], Writing—original draft [lead], Writing—review & editing [lead])

Ting Zhao (Conceptualization [lead], Supervision [lead], Validation [lead], Visualization [lead], Writing—review & editing [lead])

## Acknowledgement

We thank Professor Lin Li for helpful discussions on RNA-seq analysis on signal transduction.

## Supplement Materials

See supplement zip files for all tests results and the statics distributions.

