## Supplemental Matrial for "resolveS: rapid inference of RNA-seq library strandedness using universal rRNA alignments": s1_data_clean.R

```
rm(list = ls())
# s1_data_clean.R
library(tidyverse)

# 设置输入目录
input_dir <- "./input_stats_files/20260104"
meta_dir <- "./input_stats_files/20251226"  # metadata 用原来的

# ============================================================================
# 1. Naphade_et_al_2022
# ============================================================================
message("Processing Naphade_et_al_2022...")

# 读取 pair 结果文件
naphade <- read_tsv(file.path(input_dir, "Naphade_et_al_2022", "output.results.01032050.tsv"),
                    show_col_types = FALSE)

# 读取 metadata
naphade_meta <- read_tsv(file.path(meta_dir, "Naphade_et_al_2022", "SRP308256_meta2.tsv"),
                         show_col_types = FALSE) %>%
  select(run_accession, experiment_title) %>%
  mutate(sample_name = str_extract(experiment_title, "(?<=: )[^:]+(?= Homo sapiens)")) %>%
  select(run_accession, sample_name)

# 从 File 字段提取 SRR 编号
naphade <- naphade %>%
  mutate(run_accession = str_extract(index_str, "SRR\\d+"))

# 合并 metadata
df_clean_Naphade_et_al_2022 <- naphade %>%
  left_join(naphade_meta, by = "run_accession")

message(paste("Naphade_et_al_2022:", nrow(df_clean_Naphade_et_al_2022), "rows"))

# ============================================================================
# 2. self_data
# ============================================================================
message("Processing self_data...")

# 读取 pair 结果文件
self_data <- read_tsv(file.path(input_dir, "Self_Data", "output.results.01032050.tsv"),
                      show_col_types = FALSE)

# 从 File 提取信息
df_clean_self_data <- self_data %>%
  mutate(Experiment = str_extract(index_str, "(?<=align\\.).*(?=\\.default\\.sam)"))

message(paste("self_data:", nrow(df_clean_self_data), "rows"))

# ============================================================================
# 3. Signal_2022
# ============================================================================
message("Processing Signal_2022...")

# 读取 pair 结果文件
signal <- read_tsv(file.path(input_dir, "Signal_2022", "output.Signal_2022.results.01032050tsv"),
                   show_col_types = FALSE)

# 读取 metadata（已筛选最大采样数的版本）
signal_meta <- read_tsv(file.path(input_dir, "Signal_2022", "12859_2022_4572_MOESM2_ESM_max_sampled_resolveS_filtered.tsv"),
                        show_col_types = FALSE,
                        comment = "##")

# 获取最后两列的列名
last_two_cols <- names(signal_meta)[(ncol(signal_meta) - 1):ncol(signal_meta)]

# 选取需要的列（StrandedenessType 改用 Caution 列）
signal_meta_clean <- signal_meta %>%
  select(
    Species,
    Sample,
    StrandedenessType = Caution,
    StrandedenessJudge = all_of(last_two_cols[1]),
    JudgeStatus = all_of(last_two_cols[2])
  )

# 从 File 提取 Sample（SRR 编号）
signal <- signal %>%
  mutate(Sample = str_extract(index_str, "SRR\\d+"))

# 合并 metadata
df_clean_Signal_2022 <- signal %>%
  inner_join(signal_meta_clean, by = "Sample")

message(paste("Signal_2022:", nrow(df_clean_Signal_2022), "rows"))

# "h. sapiens"    "s. cerevisiae" "a. thaliana"
df_clean_Signal_2022 %>% filter(Species == "s. cerevisiae") %>%
  select(Sample, Strandedness, NeedPrecise, Fwd, Rev, Fwd_Ratio,Rev_Ratio,Rel_Diff , StrandedenessType, Species) %>% View()

df_clean_Signal_2022 %>% filter(Species == "h. sapiens") %>%
  select(Sample, Strandedness, NeedPrecise, Fwd, Rev, Fwd_Ratio,Rev_Ratio,Rel_Diff , StrandedenessType, Species) %>% View()

df_clean_Signal_2022 %>% filter(Species == "a. thaliana") %>%
  select(Sample, Strandedness, NeedPrecise, Fwd, Rev, Fwd_Ratio,Rev_Ratio,Rel_Diff , StrandedenessType, Species) %>% View()

df_clean_Signal_2022$Species %>% unique()

# write the reads to the dir
#write_tsv(df_clean_Signal_2022, file = file.path(input_dir, "Signal_2022", "merged.output.tsv"))

df_clean_Signal_2022 %>% select(Sample, strand_type, StrandedenessType, Species, MAPQ_filter, detection_level) %>%
  mutate(is_correct = if_else(
    (strand_type == "fr-unstranded" & StrandedenessType == "Unstranded") |
      (StrandedenessType == "Stranded" & strand_type != "fr-unstranded"),
    TRUE,
    FALSE
  )) %>% View()

# ============================================================================
# 4. Sigurgeirsson_et_al_2014
# ============================================================================
message("Processing Sigurgeirsson_et_al_2014...")

# 读取 pair 结果文件（制表符分隔）
sigur <- read_tsv(file.path(input_dir, "Sigurgeirsson_et_al_2014", "output.Sigurgeirsson2014.results.01032050tsv"),
                  show_col_types = FALSE)

# 读取 metadata
sigur_meta <- read_tsv(file.path(meta_dir, "Sigurgeirsson_et_al_2014", "filereport_read_run_PRJNA251691.tsv"),
                       show_col_types = FALSE) %>%
  select(run_accession, experiment_alias)

# 从 File 提取 SRR 编号
sigur <- sigur %>%
  mutate(run_accession = str_extract(index_str, "SRR\\d+"))

# 合并 metadata
df_clean_Sigurgeirsson_et_al_2014 <- sigur %>%
  left_join(sigur_meta, by = "run_accession")

message(paste("Sigurgeirsson_et_al_2014:", nrow(df_clean_Sigurgeirsson_et_al_2014), "rows"))

# ============================================================================
# 打印数据概览
# ============================================================================
message("\n========== Data Summary ==========")

message("\n--- df_clean_Naphade_et_al_2022 ---")
glimpse(df_clean_Naphade_et_al_2022)

message("\n--- df_clean_self_data ---")
glimpse(df_clean_self_data)

message("\n--- df_clean_Signal_2022 ---")
glimpse(df_clean_Signal_2022)

message("\n--- df_clean_Sigurgeirsson_et_al_2014 ---")
glimpse(df_clean_Sigurgeirsson_et_al_2014)

message("\nData cleaning completed successfully!")

# 保存为 .RData
save(df_clean_Naphade_et_al_2022,
     df_clean_self_data,
     df_clean_Signal_2022,
     df_clean_Sigurgeirsson_et_al_2014,
     file = "./input_stats_files/20260104/clean_data.RData")
```
