## Supplemental Matrial for "resolveS: rapid inference of RNA-seq library strandedness using universal rRNA alignments": s2_counting_datasets.R

```
rm(list = ls())
library(tidyverse)

sample_size <- nrow(df_clean_Naphade_et_al_2022) + nrow(df_clean_self_data) +
  nrow(df_clean_Signal_2022) + nrow(df_clean_Sigurgeirsson_et_al_2014)


error_size <- 3

error_ratio <- error_size / sample_size

1 - error_ratio
```
