## Supplementary figures and images for "resolveS: rapid inference of RNA-seq library strandedness using universal rRNA alignments"

### S1_self_data_raincloud_plots.pdf

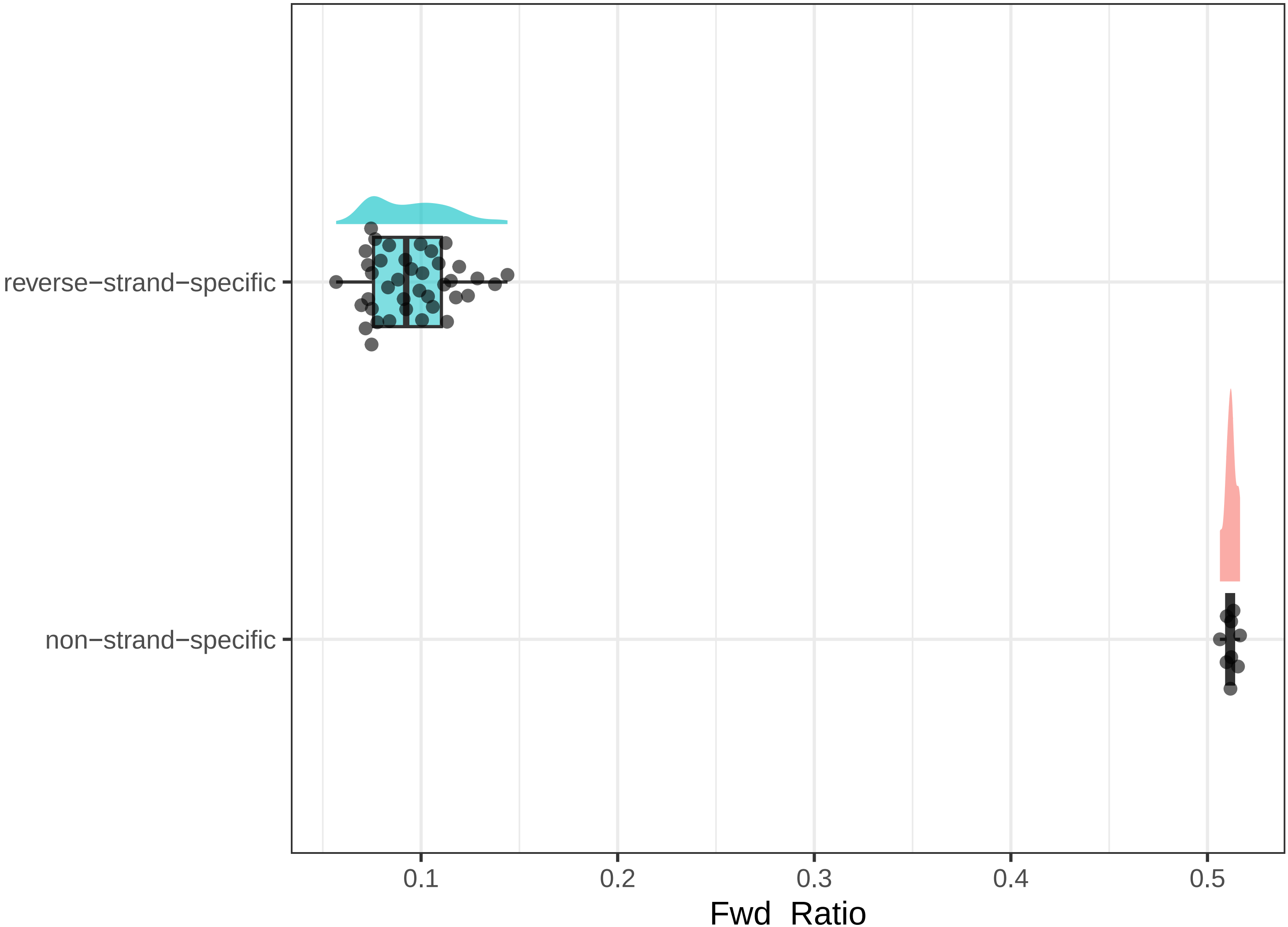

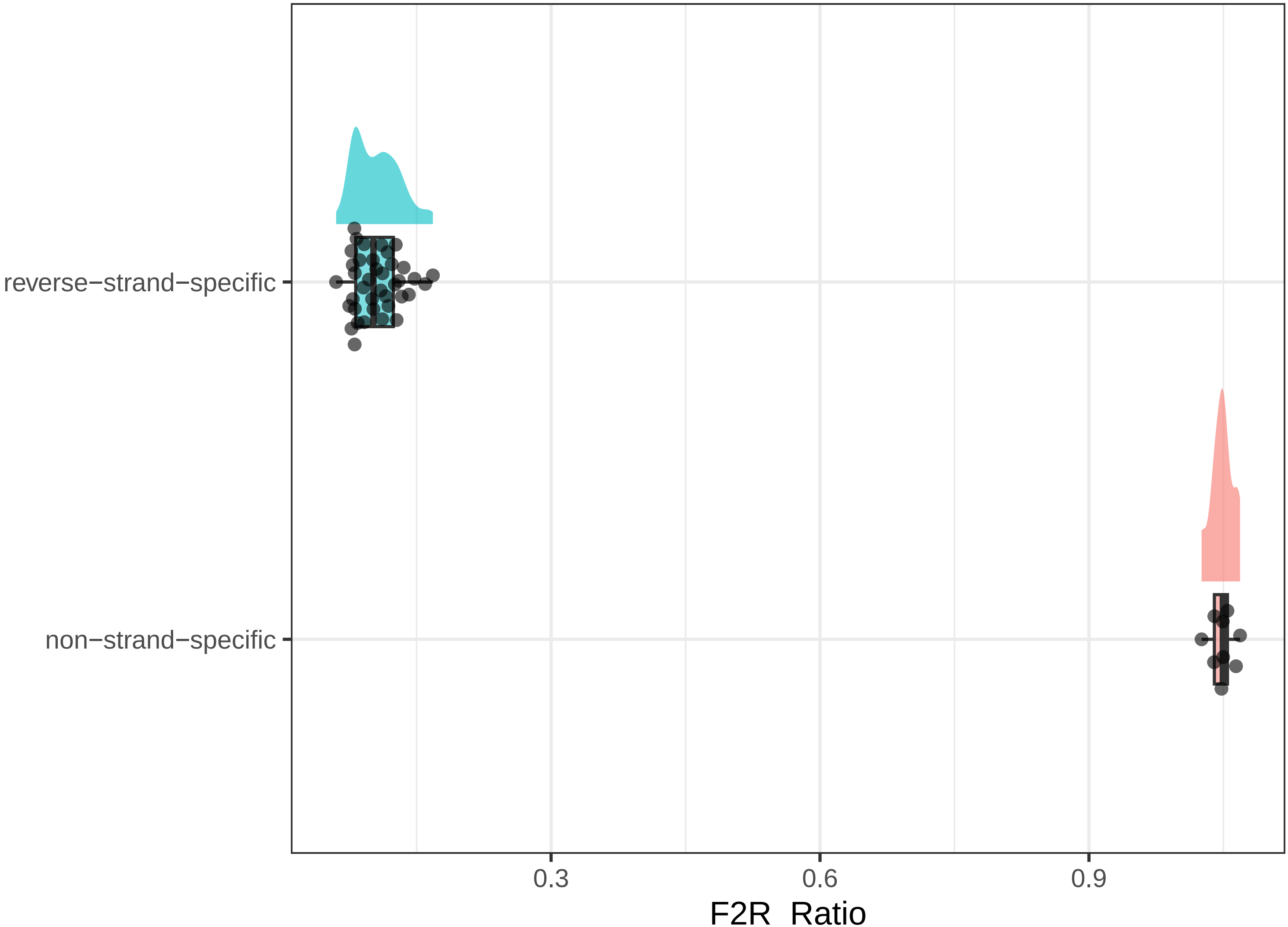

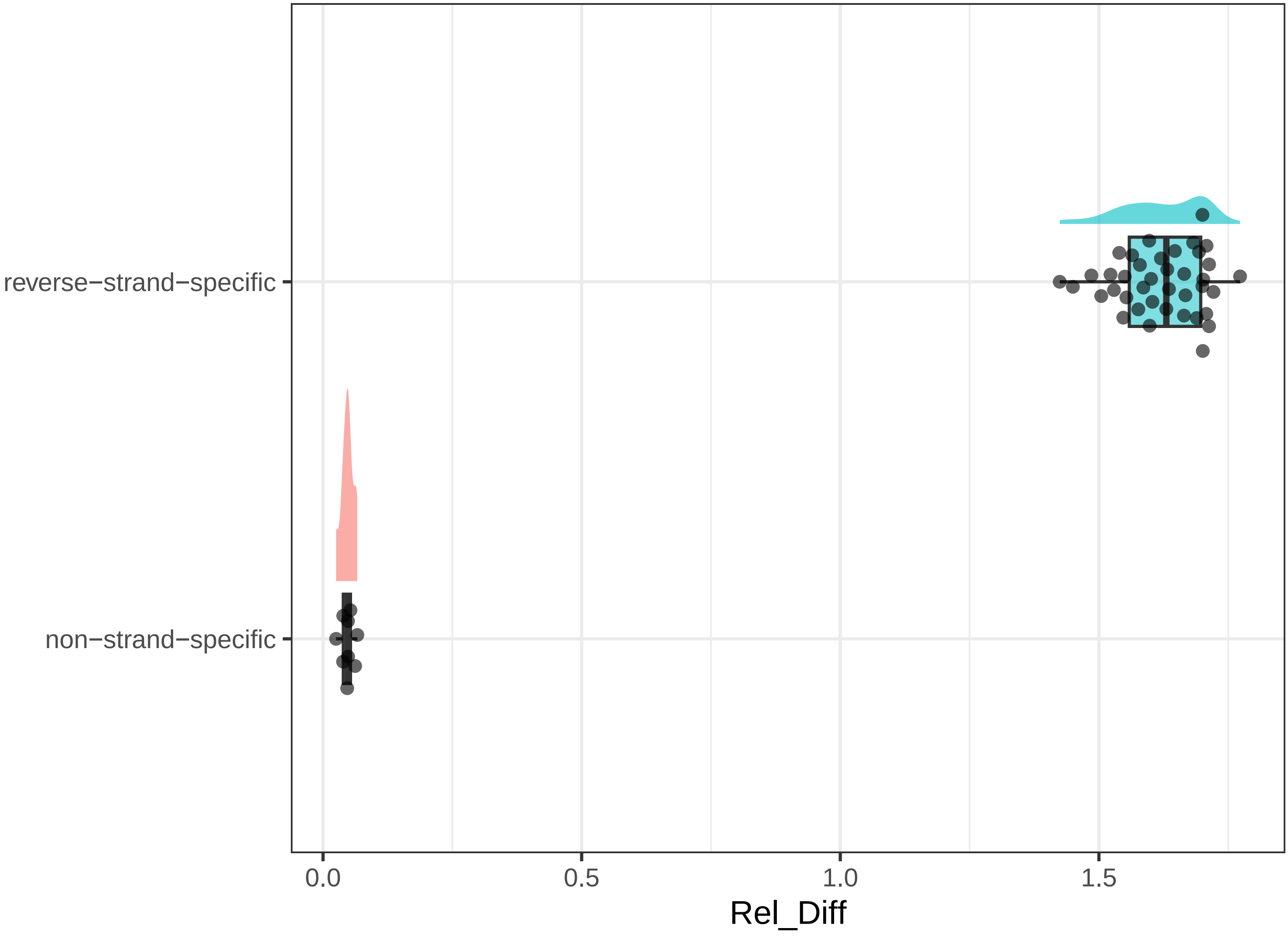

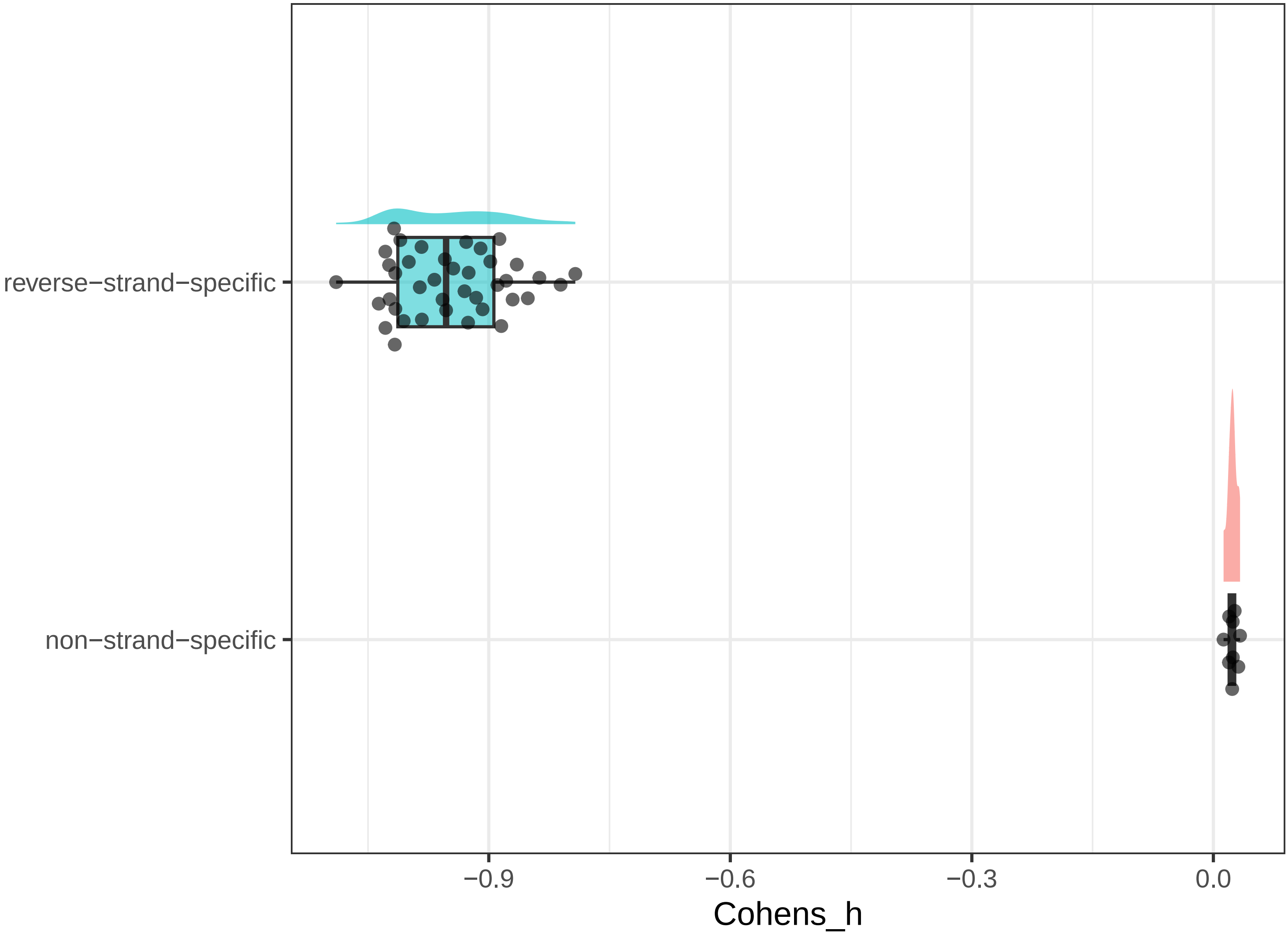

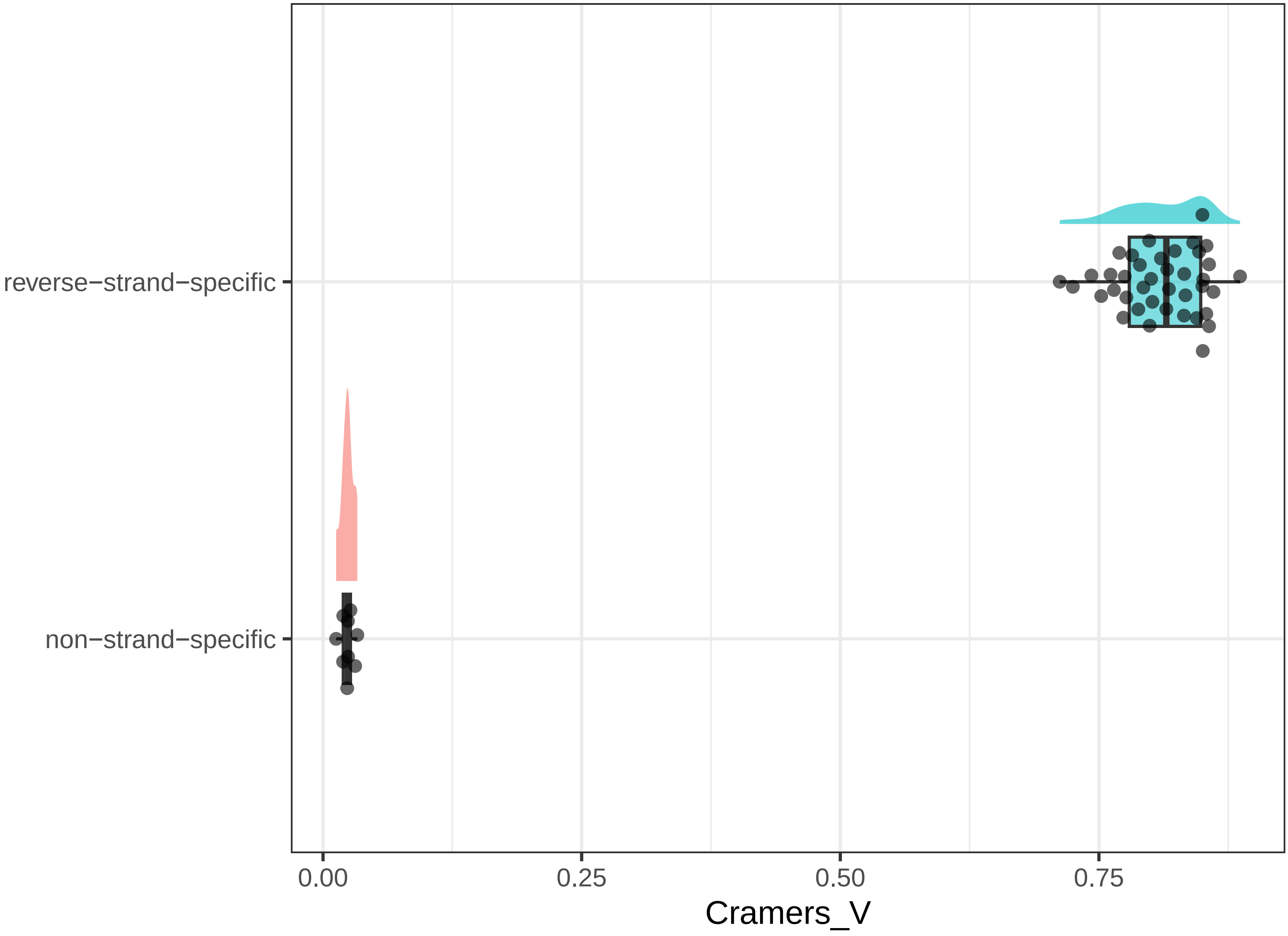

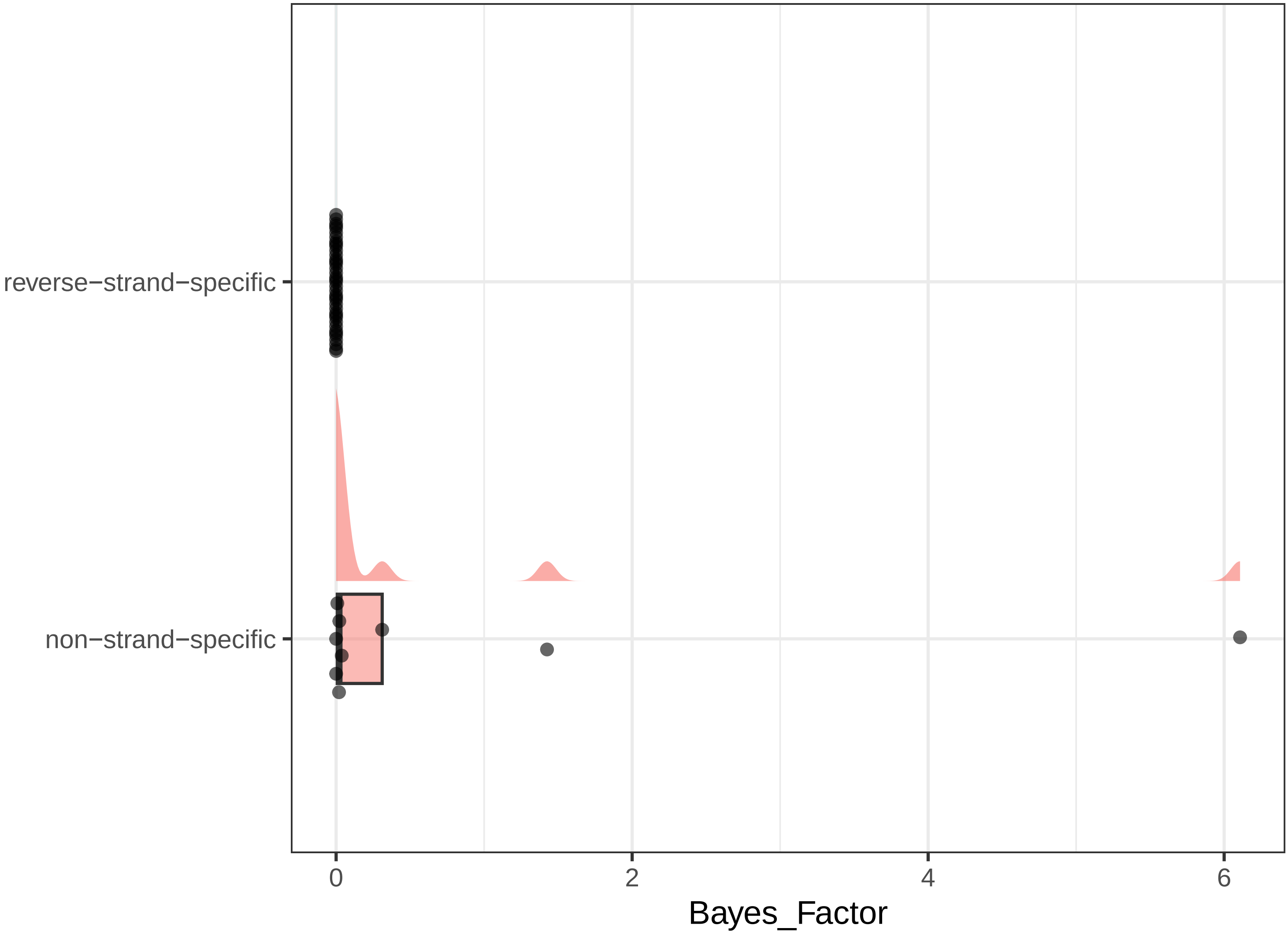

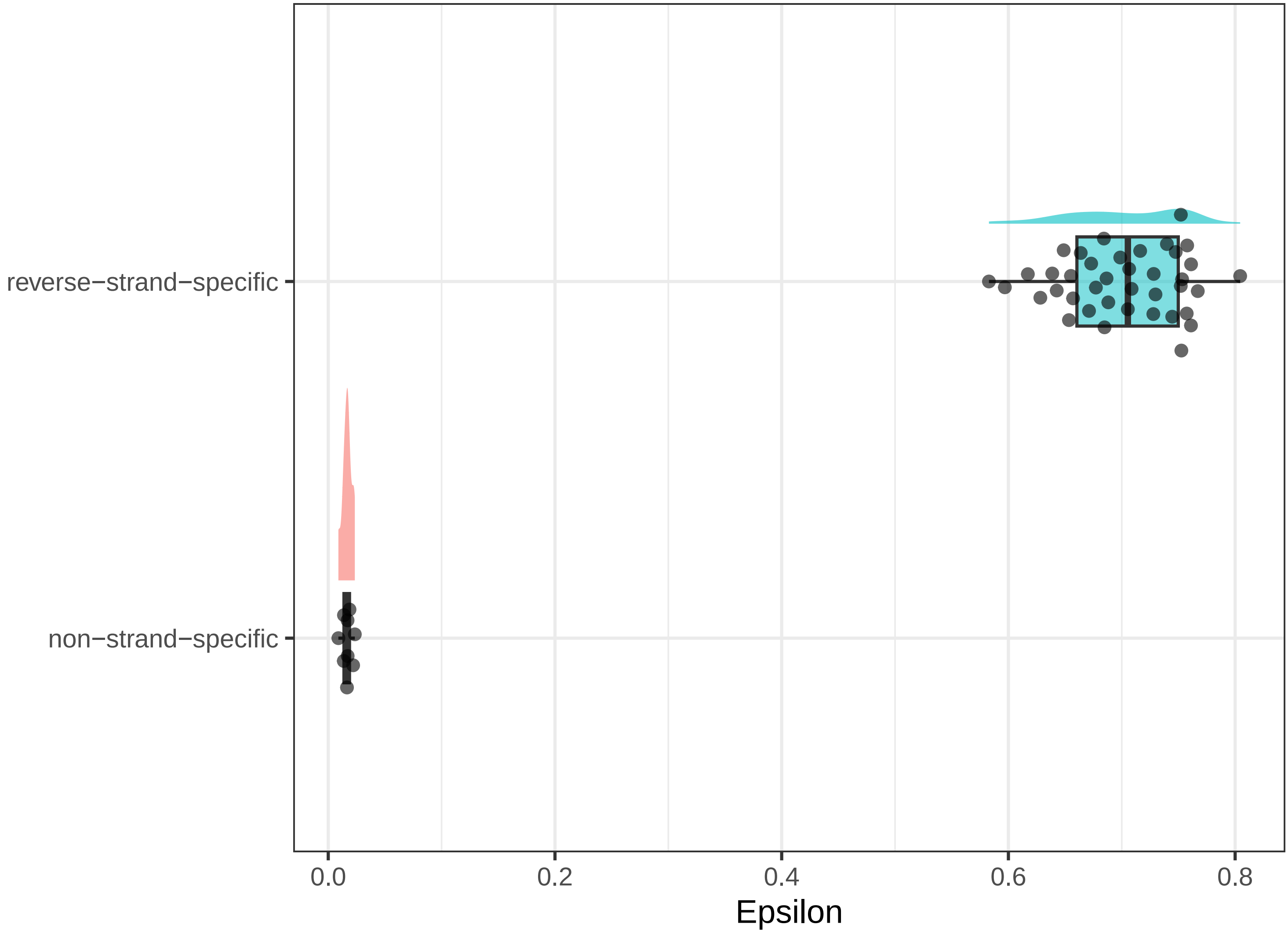

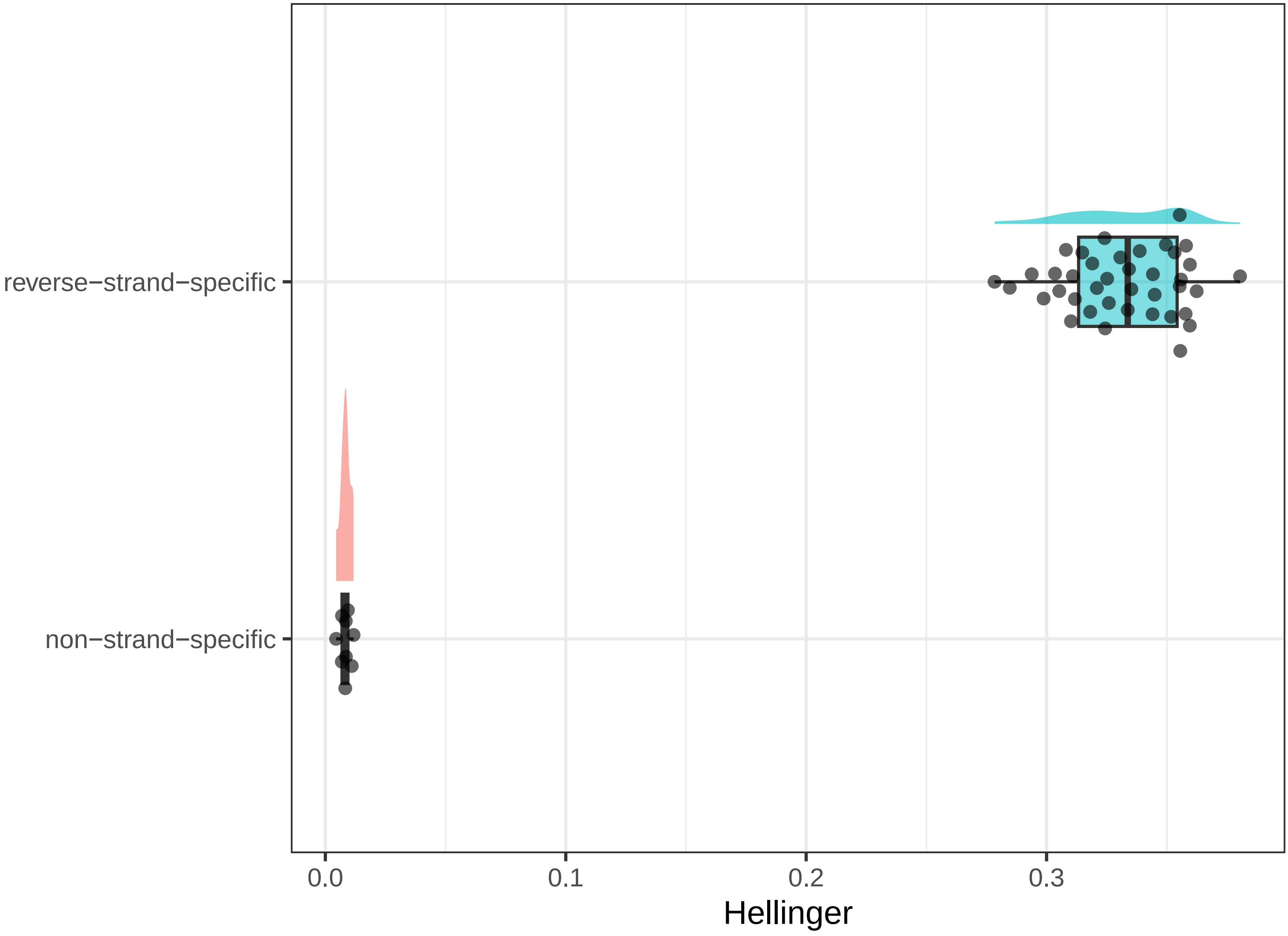

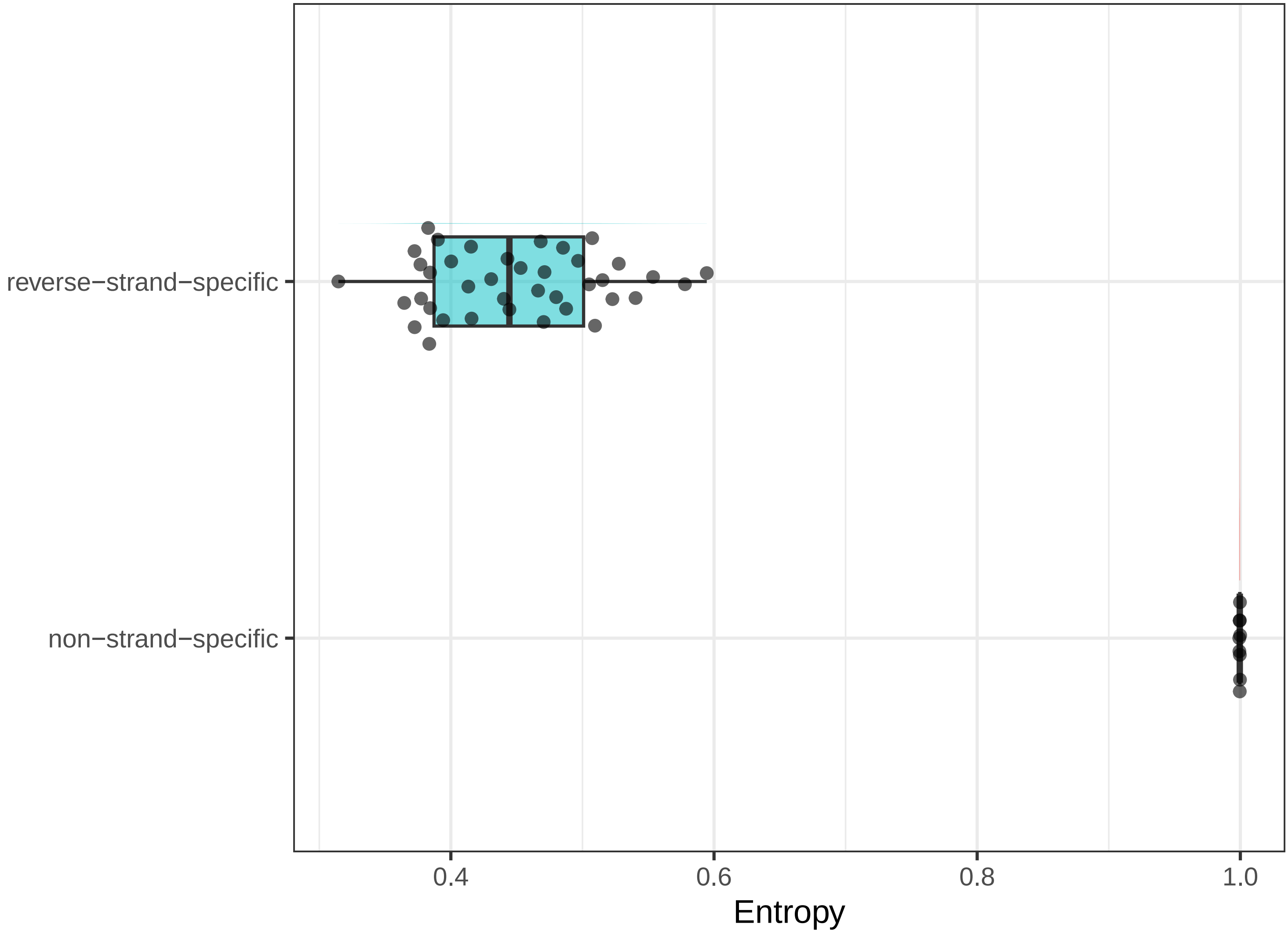

### S2_Sigurgeirsson et al., 2014_15samples_raincloud_plots.pdf

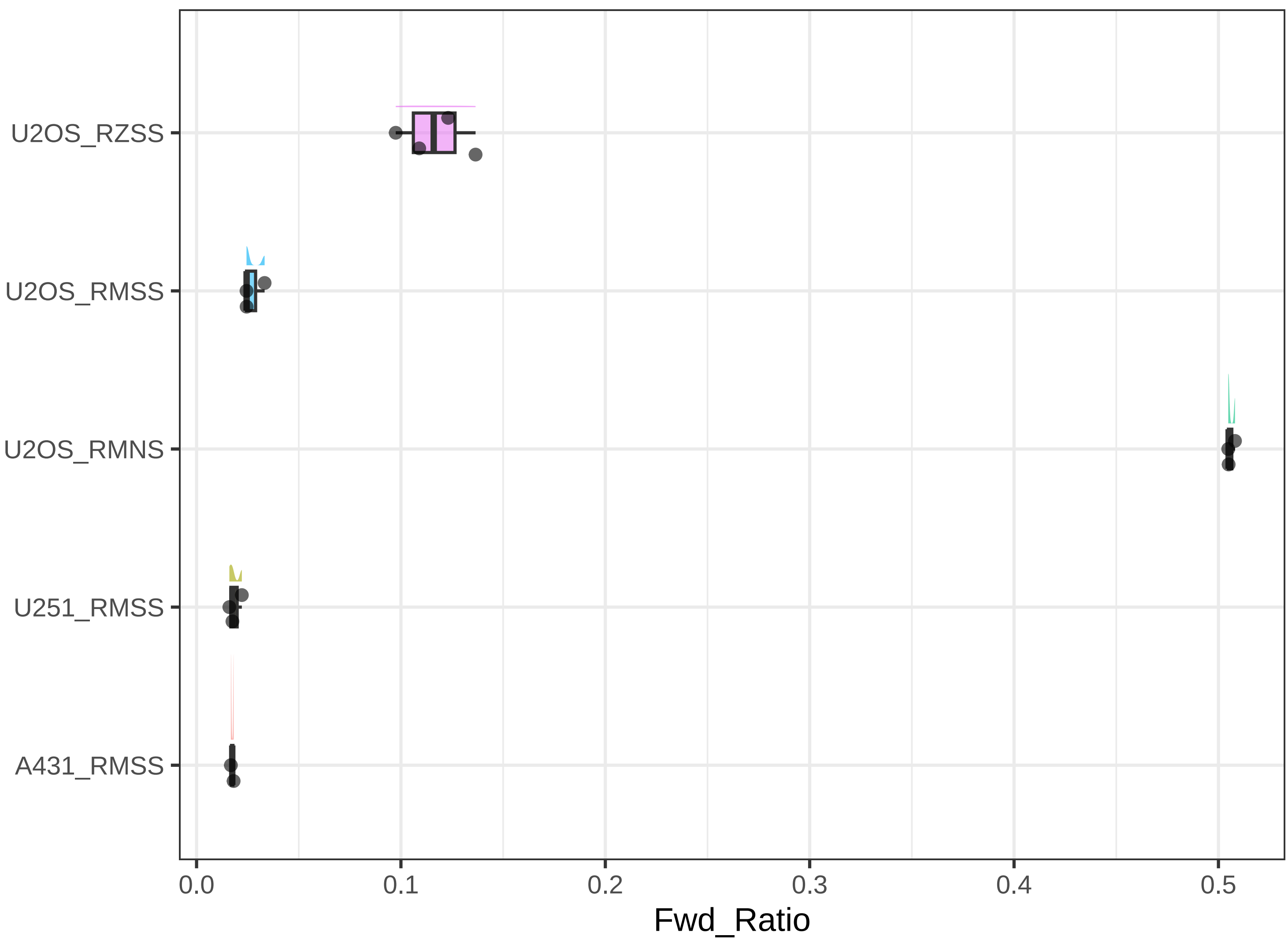

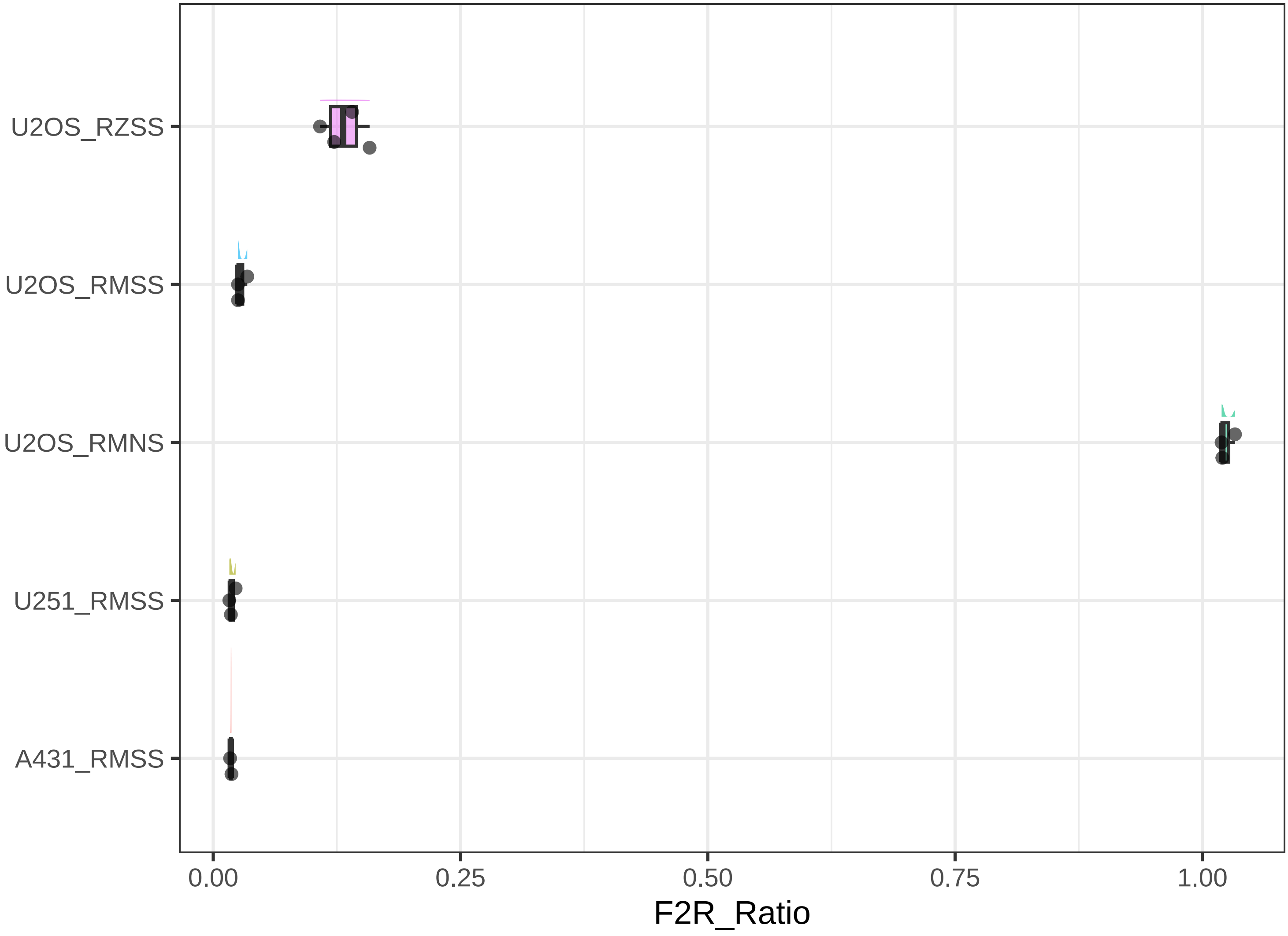

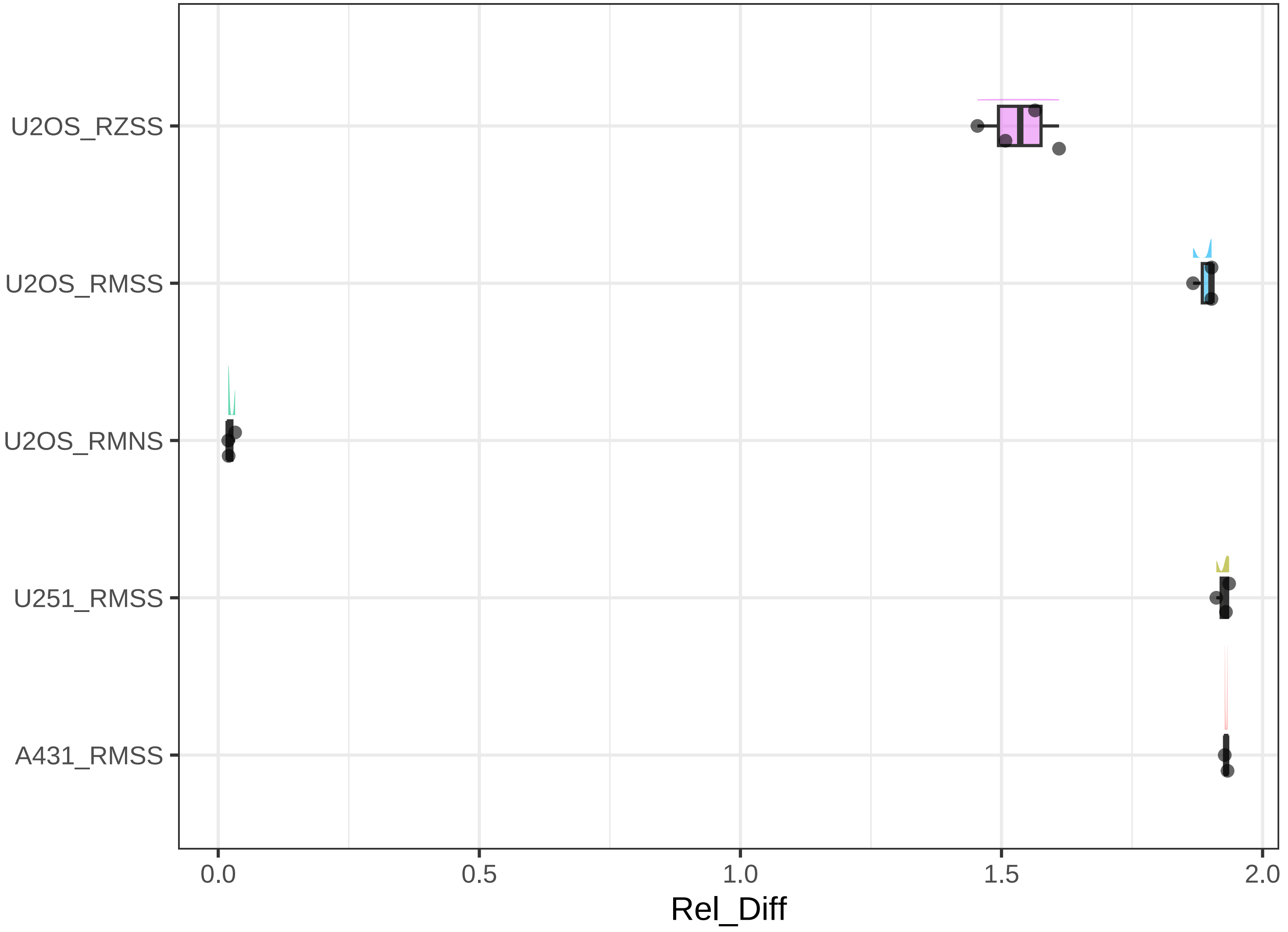

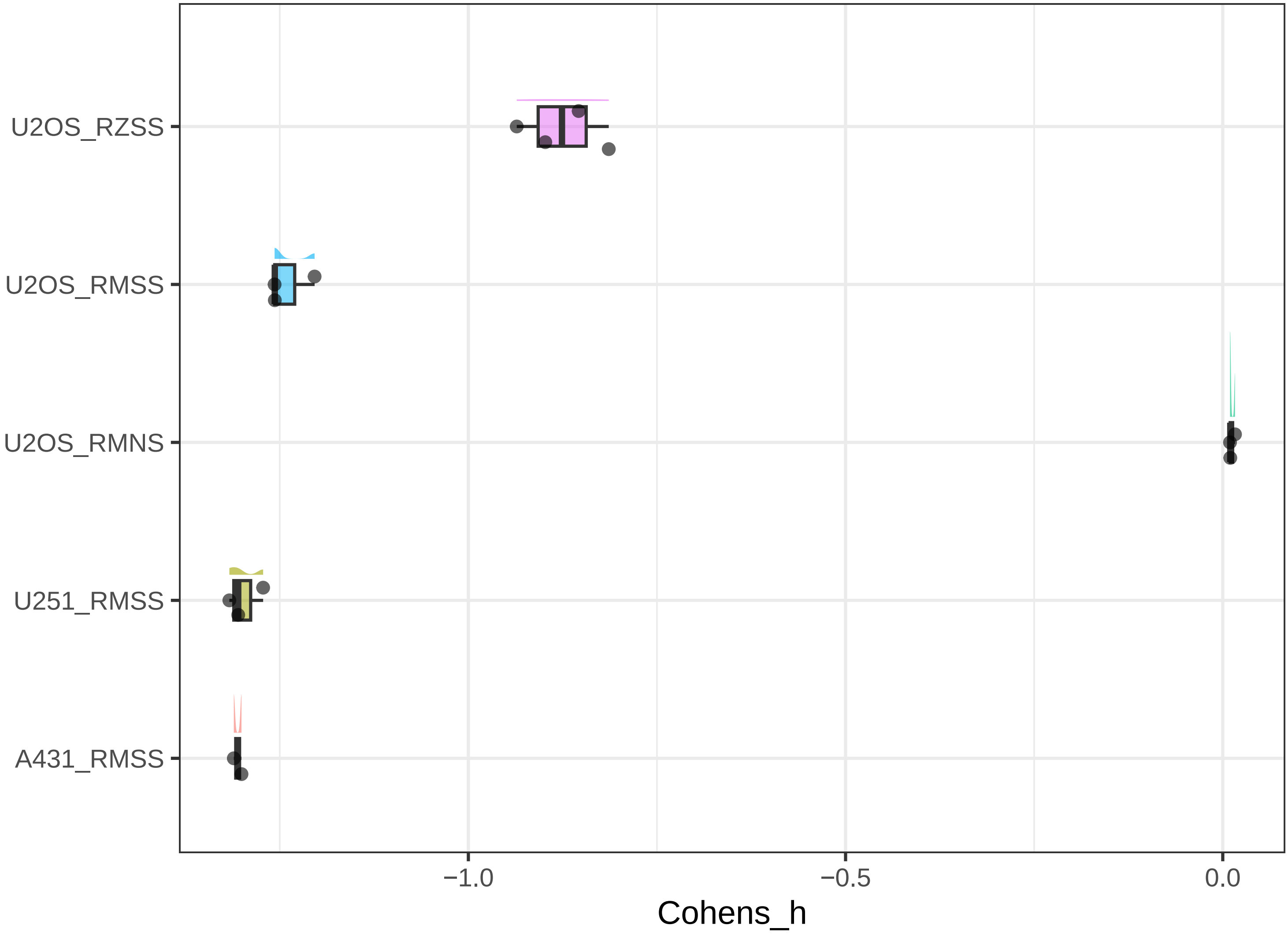

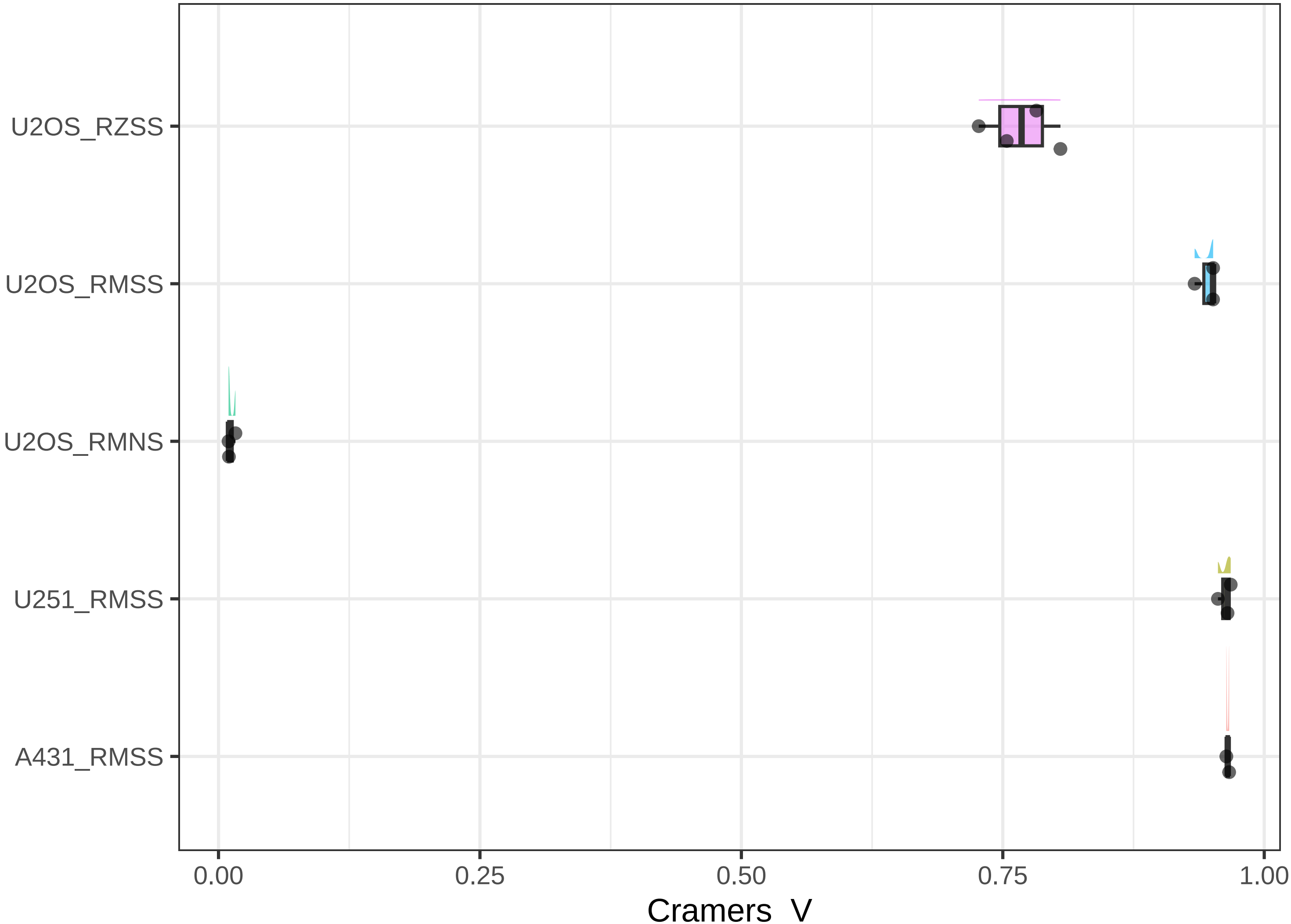

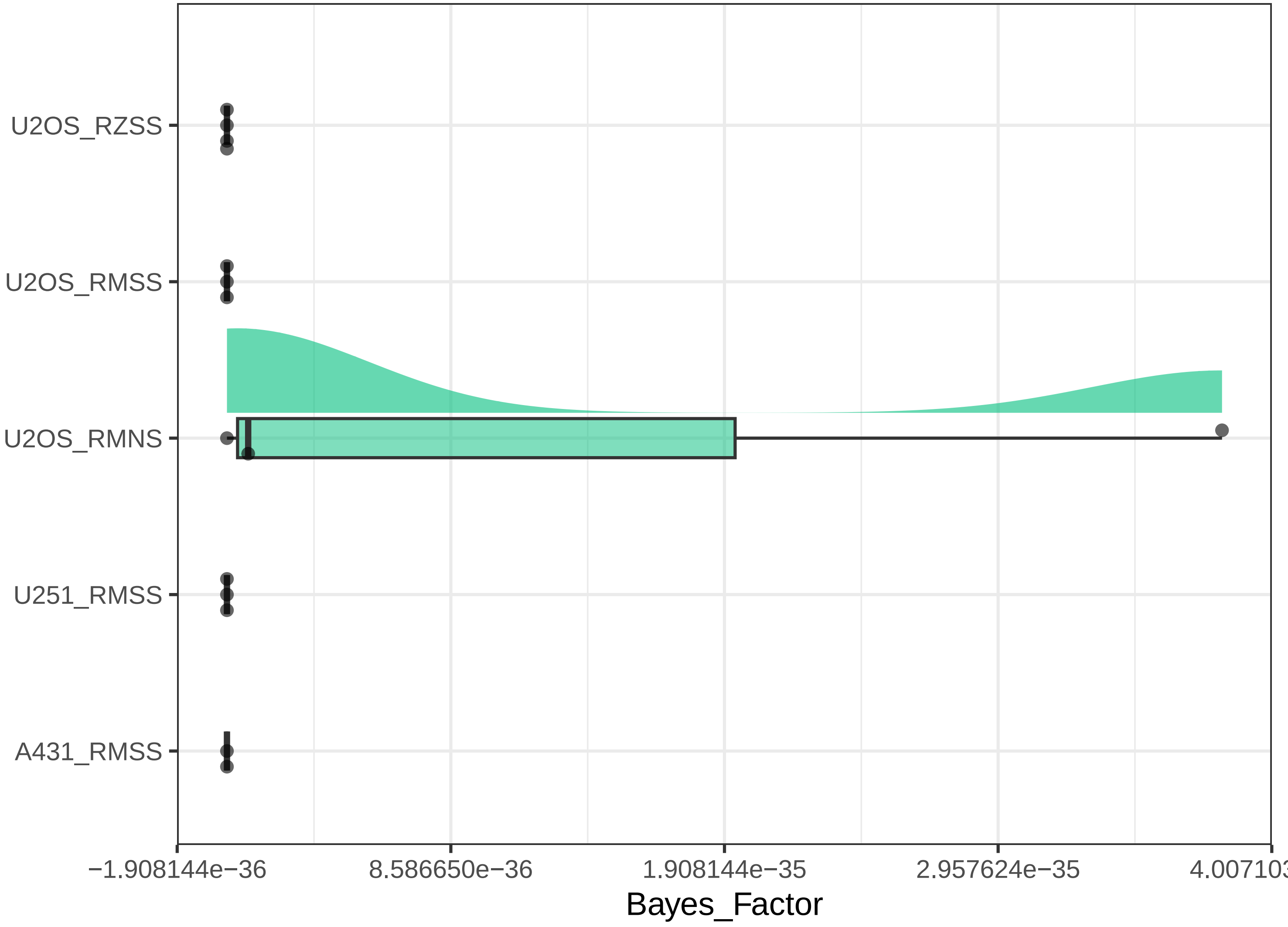

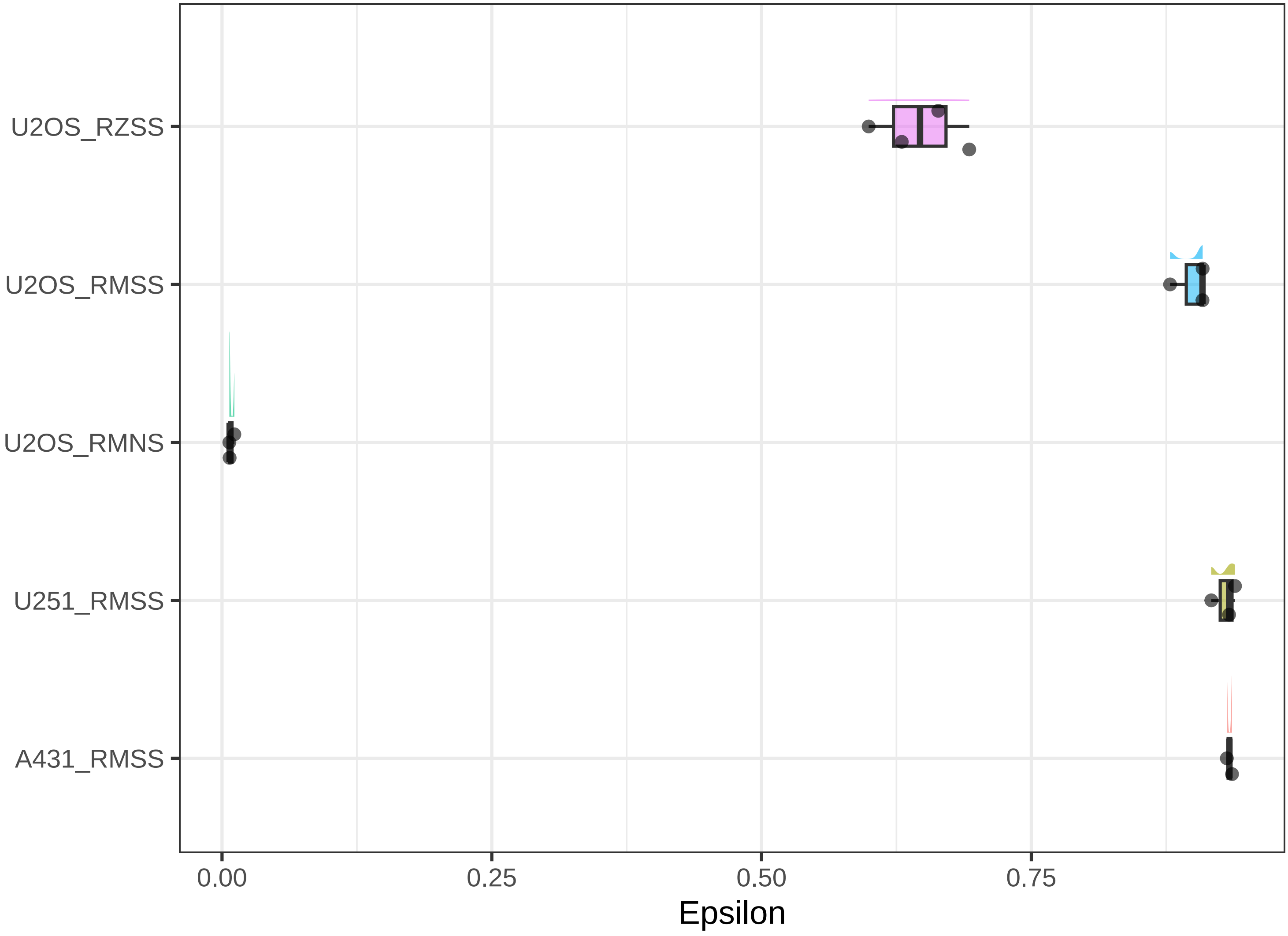

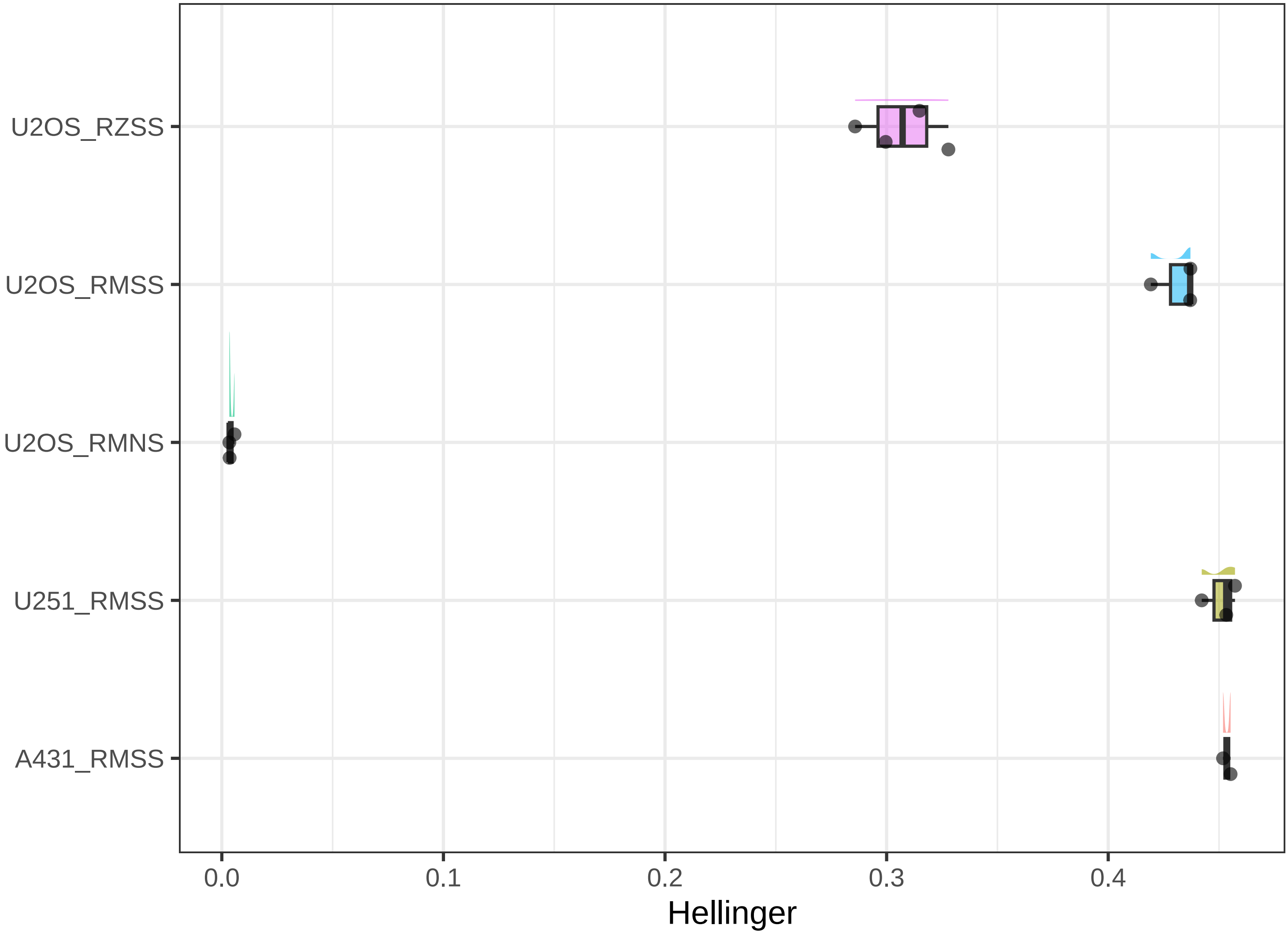

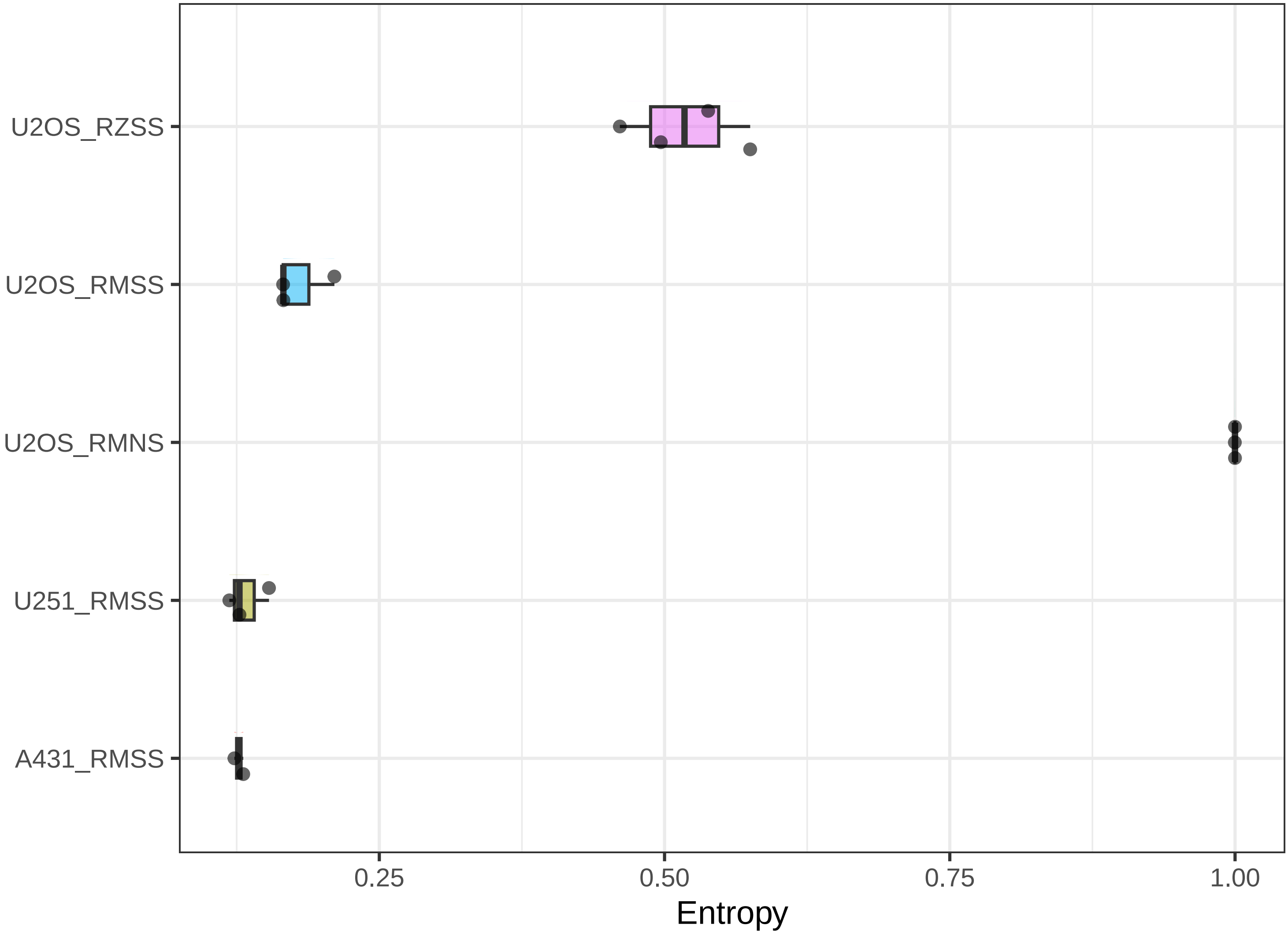
